# Coding sequence clustering universally predicts fine- and coarse-scale chromatin compartment landscapes

**DOI:** 10.64898/2026.02.26.708372

**Authors:** Rory T. Cerbus, Kyogo Kawaguchi, Ichiro Hiratani

**Affiliations:** Laboratory for Developmental Epigenetics, RIKEN Center for Biosystems Dynamics Research (BDR), 2-2-3 Minatojima-minamimachi, Chuo-ku, Kobe, 650-0047, Japan; Nonequilibrium Physics of Living Matter Laboratory, RIKEN Pioneering Research Institute, 2-1 Hirosawa, Wako, 351-0198, Japan; Institute for Physics of Intelligence, Department of Physics, The University of Tokyo, 7-3-1 Hongo, Bunkyo-ku, Tokyo, Japan; Universal Biology Institute, The University of Tokyo, 7-3-1 Hongo, Bunkyo-ku, Tokyo, Japan

## Abstract

Intra-chromosomal contact maps from many species display a striking plaid pattern, reflecting chromatin compartmentalization at a sub-chromosomal scale. Although widely regarded as a core feature of genome architecture, such patterns are absent in many organisms, and their underlying determinants remain unclear. Here we systematically examine the relationships among sequence features, chromatin compartmentalization, and evolutionary conservation across 247 species spanning five kingdoms. By testing the usual genomic suspects as determinants of compartmentalization within and between species, we identify the coding DNA sequence (CDS) density landscape as the most consistent predictor of compartmentalization, including whether an organism exhibits a fine-scale plaid or coarse-scale non-plaid chromatin contact map. In contrast, correlations with GC content, CpG density, or repeat element composition vary across lineages, contextualizing long-standing observations such as the prominence of chromosome G-banding in amniotes and its absence elsewhere. Notably, compartment organization is conserved across syntenic blocks between species separated by up to 1 billion years of evolution, and this conservation tracks with preservation of CDS density profiles rather than other sequence features. These findings establish the genomic distribution of coding sequences as a universal and deeply conserved organizing principle of nuclear compartment architecture.

## Introduction

Eukaryote genomes are compacted and organized within the cell nucleus. Some of the earliest experimental approaches to provide a visual indication of the complexity of this three-dimensional (3D) organization were chromosome banding techniques [1, 2, 3]. The mosaic patterns they revealed, at least in amniotes, hinted at a complex yet highly ordered genome architecture that is specific to each species. The 3D structure of the genome can now be probed more directly by techniques such as Hi-C (high-throughput chromosome conformation capture) technology [4], which determines the contact frequency between different genomic regions to generate genome-wide chromatin contact maps. Examination of these maps across scales reveals a hierarchical organization, ranging from kilobase-scale chromatin loops [5] to sub-gigabase (Gb) inter-chromosomal contact patterns [6].

One of the first structures to be elucidated by Hi-C was the A/B chromatin compartment(s) [4], hereafter referred to simply as compartment(s) for brevity. When viewing many Hi-C contact maps at the chromosome scale, a striking plaid pattern leaps into view, revealing large domains that are insulated from nearby domains yet preferentially interact with distant ones (Fig. 1a, Hi-C contact map). This checkered pattern can be decomposed into at least two juxtaposed compartments, which are typically ∼0.04 times the chromosome size, or on the order of several megabases (Mb) in mouse and human (see Extended Data Fig. 1a), and reminiscent of classical chromosome banding patterns. However, their genomic determinants and the mechanisms underlying their formation remain largely unresolved [7, 8, 9], in stark contrast to smaller-scale structures such as topologically associating domains (TADs) [10, 11, 12, 13, 14] and even larger, inter-chromosomal contact patterns [6].

**Fig. 1.**
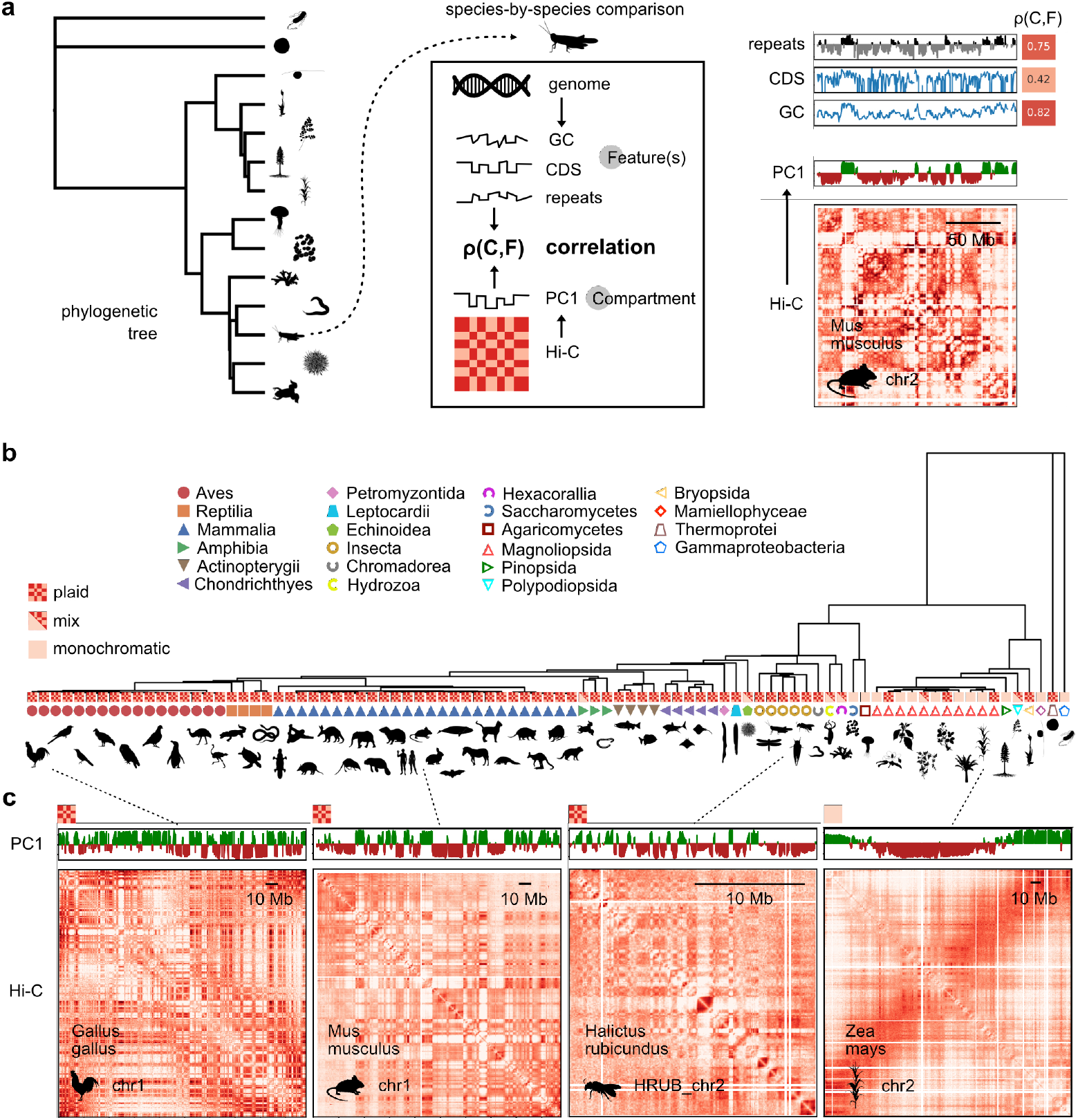
Comparing the chromatin compartments across the tree of life. (a) Schematic of our approach. Genomic and Hi-C data are sampled from organisms across the tree of life. The compartments (*C*) are extracted as the first Principal Component (PC1), which is compared against various genomic features (*F*). We focus on the Pearson correlation, ρ(*C, F*). (b) Phylogenetic tree at the order level showing the plaid/monochromatic trait at the tip. Some branches, labeled “mix”, had both traits exhibited by different chromosomes. (c) Examples of chromosome-scale Hi-C contact maps and the corresponding Hi-C PC1 for chicken (*G. gallus*), mouse (*M. musculus*), orange-legged furrow bee (*H. rubicundus*), and corn (maize; *Z. mays*) from left to right. The contact maps of most species investigated reveal the characteristic plaid pattern which is associated with a typical chromatin compartment organization. Corn and some other species do not exhibit a plaid pattern, and the corresponding PC1 is instead a large-scale variation mode.

A substantial set of the instructions for compartmentalization appears to be encoded in the genome sequence. Notwithstanding cell type differences (see Extended Data Figs. 1c), numerous studies on mice and humans, and fewer studies on other mammals, have shown that the euchromatic or A compartments are gene-rich [4], exhibit higher GC content [15], and are enriched for short (B1/Alu) repeat elements [16], a class of short interspersed nuclear elements (SINEs). The heterochromatic or B compartments, by comparison, are gene- and GC-poor and show higher densities of long (L1) repeats [16], a class of long interspersed nuclear elements (LINEs). More recently, computational models developed for human and mouse have suggested that the rules for not only the compartments but even the Hi-C contact maps themselves are largely encoded in the genome sequence [17, 18].

Another prominent approach to identify conserved genomic instructions is to perform pairwise comparisons of genomic sequence and compartmentalization in parallel. Imakaev et al. reported that compartment scores on 1-Mb blocks of syntenic sequence between human and mouse were strongly correlated, and yet this correlation could not be fully explained by similarities in local GC content [19]. When the binarized A/B compartment profiles within syntenic blocks were compared between four mammalian species, Álvarez et al. found broadly similar compartmentalization patterns, although TAD organization was even more strongly conserved across these same species [20, 21]. Comparing compartment values and insulation scores at finer resolution (≃10 kb) across homologous bins in 10 mammalian species, Li et al. also found phylogeny-dependent similarity in these segmented compartments but stronger conservation for insulation [22]. Comparing A/B compartmentalization across syntenic blocks from 11 mammalian species within Carnivora, Corbo et al. [23] found striking visual resemblance for sizes up to chromosomal scale; notably, they even found that syntenic blocks in shuffled genomes visually maintained similar compartmentalization patterns, suggesting that conservation of chromatin organization is linked to sequence conservation.

Taken together, these studies support the view that many of the rules governing canonical A/B compartmentalization are encoded in the genome sequence, of which specific candidate sequence features have been proposed in mouse, human, and other model organisms [24]. Although it is tempting to assume that these relationships generalize to other regions of the tree of life, large-scale studies of compartmentalization have never been carried out. As will be discussed below, many species do not even exhibit the typical plaid pattern characteristic of mammalian A/B compartmentalization, or only some of their chromosomes do, suggesting the possibility of alternative organizational principles. Here we compared genome sequences and Hi-C contact maps from 247 species spanning 162 families, 89 orders, including 26 of 27 mammalian orders, 22 classes, 14 phyla, and 5 kingdoms, to identify general rules of compartmentalization.

## Results

### Hi-C analysis of 247 species reveals plaid and monochromatic Hi-C contact maps

To identify the rule(s) encoded in the genome sequence that govern A/B compartmentalization, we created a database of chromosome-level genome assemblies and corresponding Hi-C data mainly from NCBI [25], DNA Zoo [26, 27], and GenomeArk [28] (Fig. 1). Genomes hosted on DNA Zoo often have Hi-C contact map data already aligned to the genome assembly (e.g., orange-legged furrow bee, *H. rubicundus*; see Fig. 1c). In all other cases we aligned the raw Hi-C reads to the genome assemblies using the Juicer platform [27] with default parameters, similar to DNA Zoo. Genome assemblies without annotations or with poor BUSCO scores [29] were annotated using GeMoMa (v1.9) [30] and Liftoff (v1.6.3) [31]. We also generated *de novo* repeat libraries for all 247 species using RepeatModeler (v2.0.6) [32] and determined the repeat locations using RepeatMasker (v4.1.8) [33]. All data were organized according to a phylogenetic tree [34] (Fig. 1a), which guided and informed subsequent analysis (see Methods for more details about analyzing genomic and Hi-C data).

In many organisms, the Hi-C contact maps exhibited the familiar plaid pattern which reflects the separation of A and B compartments. Curiously, however, there are some organisms whose Hi-C contact maps do not display a prominent plaid pattern (Figs. 1b–c). Much like the weak or absent chromosome banding in many organisms [1, 35, 36, 37], the typical approach used to extract compartments instead revealed large variations in chromosome-scale contact frequency (Fig. 1c). While the plaid pattern exists in archaea [38], they are apparently absent in bacteria [39], creating a puzzling mystery as to the phylogenetic origin of either architectural preference. Because of the relative absence of structure in the non-plaid Hi-C maps, and as a historical nod to the chromosome staining experiments of the past, we refer to these organisms as monochromatic (e.g., maize in Fig. 1c) and the organisms with a typical (mammalian) checkered contact map as plaid (e.g., chicken, mouse, and orange-legged furrow bee in Fig. 1c) in the discussion that follows. To identify which species and orders showed predominantly plaid or monochromatic behavior, we set a threshold based on the Hi-C domain size relative to the chromosome size (Ext. Data Fig. 2a) and found significant phylogenetic diversity of the plaid/monochromatic character throughout the tree of life (Fig. 1b, Ext. Data Figs. 2b–c). While this monochromatism has been noticed in plants like maize (*Z. mays*) [40] or *Arabidopsis* [41], focus was often placed on alternative techniques for extracting small-scale compartments analogous to those observed in mammals. The apparent absence of plaid compartmentalization, or at a minimum its substantial diminishment, challenged the idea of a universal architecture and further motivated our search for an explanation for this clear difference. Starting from known results in species with plaid patterns, we sought to understand whether the absence of such a pattern in other species is the result of entirely different rules for compartmentalization or not.

### Comparing the genomic “usual suspects” with A/B compartment organization

#### Phylogeny-guided comparison

We extracted candidate genomic features from chromosome-level genome assemblies, and used the ‘Iterative Correction and Eigenvector decomposition (ICE)’-corrected [19] Hi-C contact maps to estimate the compartments using the first component (PC1) from a Principal Component Analysis (PCA) of the observed-over-expected contacts (Fig. 1a), which emphasizes off-diagonal contacts. While this is a standard protocol for estimating A/B compartments, we note that there are isolated cases when this does not correspond well with the plaid pattern [4]. We compared our compartments with the estimate from cooltools (v0.7.1) [42] and found no substantial differences (see Extended Data Fig. 3). We determined the ambiguous sign of the PC1 using the coding sequence (CDS) density [4], following common practice, although we later challenged this convention in our analyses. We chose the resolution based on the genome size, typically 500 kb for eutherian mammals and 1 Mb for marsupials, such that the plaid pattern was clearly visible and so there were typically at least 100 bins for the larger chromosomes. In Extended Data Fig. 3 we demonstrate that our main correlation and length scale results are not sensitive to the resolution. We then calculated the Pearson correlation between each genomic feature track and the compartments (Fig. 1a). Although modified methods have been used to extract compartments in plants using, e.g., sub-chromosomal analysis windows [40, 41], we attempted to use the same methodology for all organisms.

The main results of our broad comparison with all taxonomic orders are represented in a Pearson correlation matrix arranged according to the phylogenetic tree in Fig. 2a, a class-median summary of correlations in Fig. 2b, and several specific examples shown in Figs. 2c–e. The heatmap in Fig. 2a includes the Pearson correlation between the compartments (*C*) and the CDS density, GC content, log-ratio of the density of all SINE to all LINE elements, and the long terminal repeats (LTR) density. We denote these correlations as ρ(*C, F*), where *C* denotes the compartments (PC1), and *F* is the genomic feature in question.

**Fig. 2.**
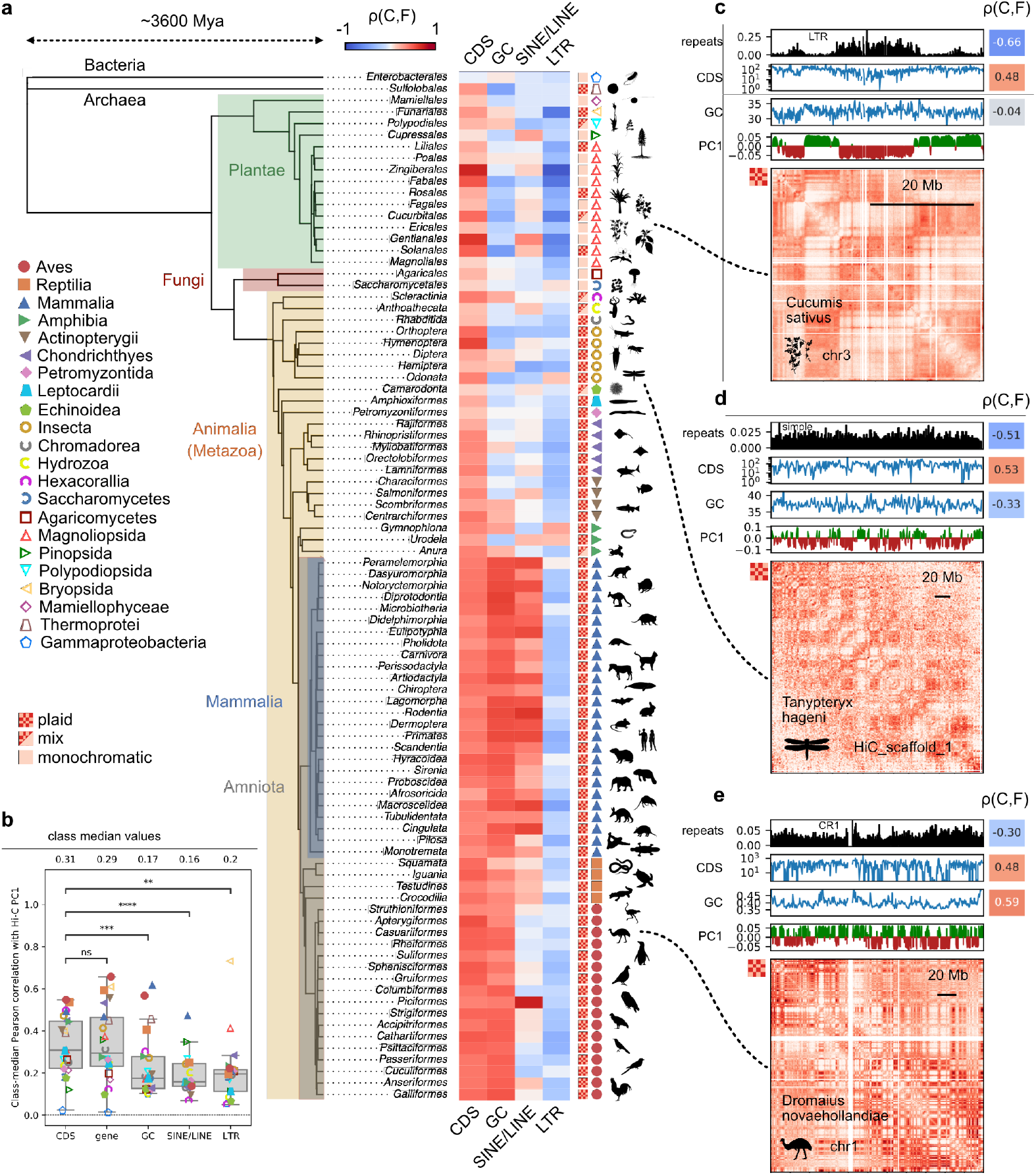
Comparison of genomic features with compartments across the tree of life. (a) A heatmap of Pearson correlations between CDS, GC, SINE/LINE and LTR densities and the Hi-C PC1 is arranged according to a phylogenetic tree at the order level. The plaid/monochromatic trait is also indicated for each order. The tree-aligned correlation heatmap indicates how various branches have specialized their relationship with compartments, such as the GC content in Amniota, the SINE/LINE density in Mammalia, and the LTR density in Plantae. (b) Distributions of the absolute values of class-median Pearson correlations where each individual genomic feature was used to independently determine the ambiguous sign of the Hi-C PC1. CDS and gene density consistently outperform other genomic features. (c)-(e) Examples of comparisons for single chromosomes from several species across the tree highlighting some of the global patterns shown on the left. Above each Hi-C contact map the Hi-C PC1 and several genomic tracks are shown. The Pearson correlation ρ(*C, F*) between each genomic feature and the Hi-C PC1 is displayed to the right of that track. (c) cucumber, *C. sativus*); (d) black petaltail dragonfly, *T. hageni*; (e) emu, *D. novaehollandiae*.

### CDS correlates with compartments in both plaid and monochromatic maps across diverse taxonomic groups

Consistent with the original findings of Lieberman-Aiden and van Berkum et al. for gene density in human lymphoblastoid cells [4], the CDS density consistently and significantly correlated with compartmentalization across orders, classes, phyla, and kingdoms, excluding bacteria, indicating a general (eukaryotic) relationship (Fig. 2a). Other well-known features such as local GC content, the logarithmic SINE/LINE density, and LTR density performed particularly well within amniotes, mammals, and plants, respectively, but outside these groups their behavior appeared erratic (Fig. 2a). Even when the PC1 sign was independently assigned using these features, thereby biasing correlations toward positive values, CDS density still stood out as an outlier (Fig. 2b). In Extended Data Fig. 1c we also show that these trends are largely independent of cell type by using compartment data from different tissues within the same species. Excluding prokaryotes (bacteria), the uniformly colored CDS column in the correlation heatmap in Fig. 2a thus signals the CDS (gene) density as a general rule for compartmentalization, in stark contrast to the non-uniform columns.

The uniformity of the correlation between PC1 and CDS density (Fig. 2a) was at first striking as well as puzzling, as this also included taxonomic orders with monochromatic Hi-C contact maps. We thus next focused on the unique genome architecture of these monochromatic species, for which the chromosomes are often trisected (or bisected) with a significant fraction of both chromosome ends being in strong contact with each other. Thus, the typical compartment size defined by PC1 could be up to nearly a third of the chromosome length (Fig. 1c, maize; Fig. 3a, subterranean clover; and Extended Data Fig. 2c). As shown in Fig. 3a, we found that the CDS density likewise displayed large-scale variations. In the center region of the chromosome of this plant (subterranean clover, *T. subterraneum*) where the PC1 value dips down, the CDS density also decays by over tenfold (Fig. 3a). Thus, for both the finer-scale plaid-patterned chromosomes and the coarsely varying ones, the PC1 of the contact map parallels the behavior of the CDS density (Fig. 3a).

**Fig. 3.**
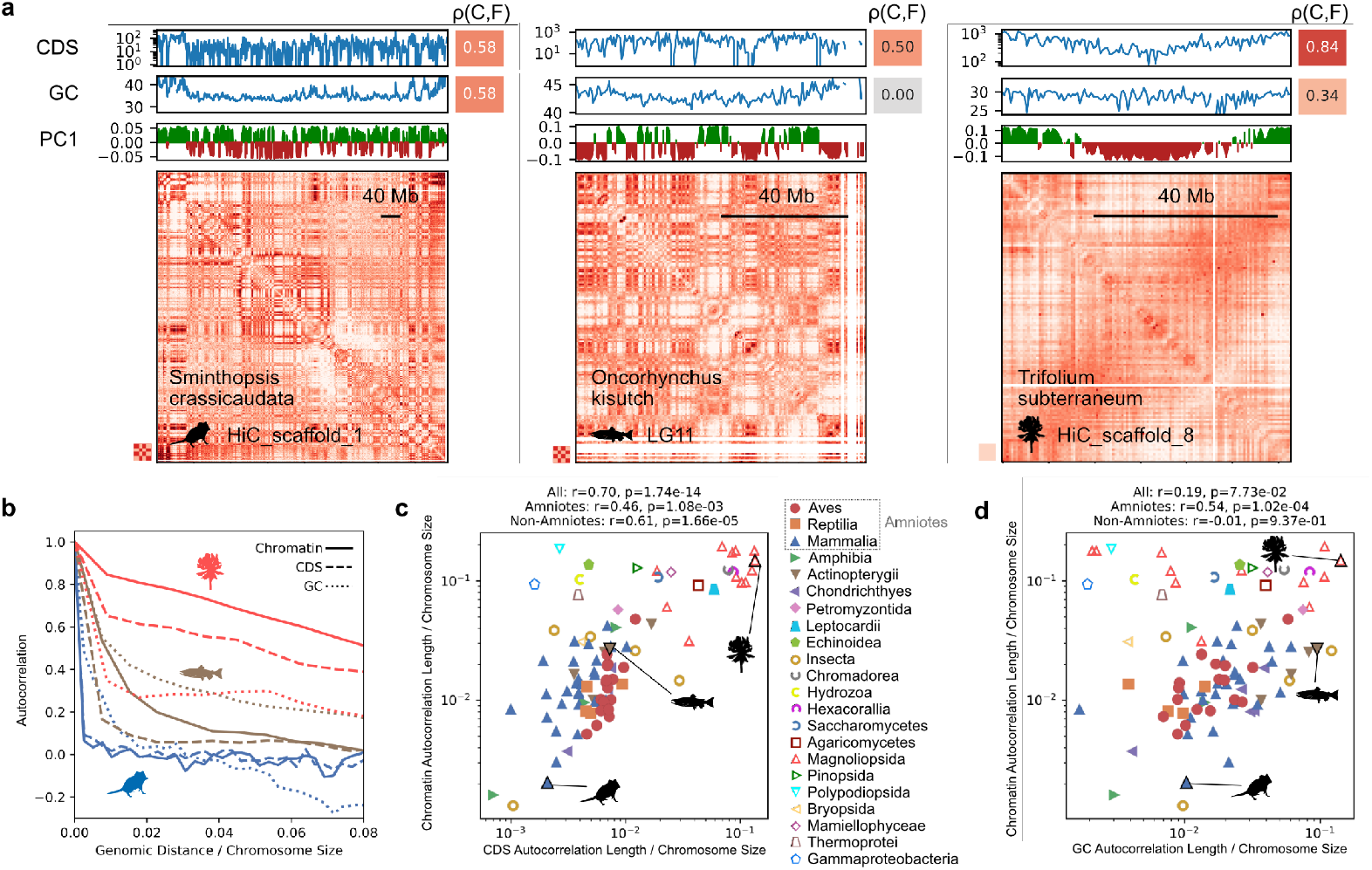
Probing the typical compartment sizes of chromatin, CDS density, and local GC content. (a) Examples of Hi-C, chromatin (A/B) compartments, local GC content, and CDS density for fat-tailed dunnart (*S. crassicaudata*), Coho salmon (*O. kisutch*), and subterranean clover (*T. subterraneum*) from left to right. The Pearson correlation *ρ*(*C, F*) between each genomic feature and the Hi-C PC1 is displayed to the right of that track. Although here the subterranean clover does not exhibit the characteristic plaid pattern found in many other species, the Hi-C PC1 nevertheless closely follows the CDS density, confirming that the relationship between higher-order chromatin structure is related to the gene density independently of the presence or absence of the plaid pattern. (b) Examples of the autocorrelation function of chromatin (Hi-C PC1), CDS density, and GC density over all sufficiently large chromosomes for these three species. All quantities decrease rapidly relative to chromosome size, except for subterranean clover which exhibits much larger-scale variations. (c,d) Order-level median plots of the characteristic size of chromatin compartments vs. the typical size of CDS density and local GC content clusters. The typical size of CDS density islands strongly correlates with the typical chromatin compartment size. We also show the Pearson correlation between these normalized correlation lengths for all orders, as well as for amniote and non-amniote orders separately.

We further probed this relationship between CDS density and Hi-C PC1 by determining characteristic length scale using the chromosome-averaged autocorrelation function (see Fig. 3b). For monochromatic species, the autocorrelation function decays slowly, reflecting the large length scales. In Fig. 3c we plotted the Hi-C PC1 characteristic length scale vs. that of the CDS density, both normalized by the average size, which indicated a strong relationship, unlike with the GC content (Fig. 3d). Extended Data Fig. 2c shows more examples of Hi-C contact maps, Hi-C PC1, and the CDS density for typical chromosomes of all the monochromatic species in our database, adding further evidence to the link between the two. Thus, the relationship between CDS density and Hi-C PC1 transcends these particulars, demonstrating its universal status.

Previous research has found a correlation between gene density and the PC1 of the Hi-C contact map in several plant species, which had chromosomes that were often trisected for gene density or Hi-C PC1 [40, 41]. By zooming in on the two ‘global’ A-compartment domains, located near the ends of each chromosome [40], they further subdivided these global domains into ‘local’ compartment domains that resemble typical compartmentalization observed in mammalian cell nuclei. Here, with our broad analysis, including 5 classes and 15 orders within the Plantae kingdom, we found that the connection between PC1 and CDS density is more general than the plaid pattern. Notably, while plants contain both plaid (e.g., cucumber; *C. sativus*), and monochromatic species, and there are also some monochromatic and mixed non-plants, the CDS to Hi-C PC1 relationship holds for all of these (Figs. 2a, 3c, Extended Data Fig. 2). The variety of heterochromatin architecture in plants has also been pointed out by Zhang et al. [43], who found that variations in heterochromatic condensates are related to variations in the sequence of the heterochromatin protein *ADCP1*. Here we found that whatever mechanism has led to the coarse chromatin structure in these plants and other species, it is also necessarily tied to the distribution of genes and CDS. We postulate that the chromosome-level gene (CDS) density is a more constrained genomic feature, involving the locations of hundreds of genes, than the behavior of a single gene or even a small group of genes [6]. Any general explanation of compartmentalization will have to account for its relationship with the heterogeneity of CDS density.

### Genomic features other than CDS density correlate with compartments only in specific taxonomic groups

While the CDS density is the most consistent indicator, other features can outperform it in specific taxonomic groups. The tight relationship between local GC content and local gene density in many species [3] would suggest that if the CDS tree is uniformly colored, then the GC content tree will be likewise. Indeed, the GC content was found early on to correlate strongly with the compartments in mammals [19] and is frequently used to determine the ambiguous sign of the principal component or eigenvector [42]. The mosaic structure of the historical banding technique appears to be linked to the presence of isochores (Mb-sized blocks of relatively homogeneous GC content) [3], thus generating the expectation that genome compartmentalization may also be tied to GC content diversity. However, outside of amniotes we found oscillations between positive and negative correlation. While most amniotes showed a strong correlation between GC content and the compartments, often stronger than between the CDS and the compartments (for instance, see Fig. 2e, emu), even some groups within amniotes, such as the order Squamata, had only weak correspondence (Fig. 2a, Extended Data Figs. 4a,d). The CpG density, which is known to correlate with methylation in some species [44], also correlated strongly within amniotes, and even tended to outperform the GC content outside this branch. And yet it also swung through positive and negative correlation, most notably in Metazoa (Animalia) (see Extended Data Figs. 4a,d,e).

Next we considered the relationship between repeat elements and compartments. Lu et al. [16] discovered that the SINE-B1/Alu SINE elements and the LINE-1 (L1) elements correlate strongly with the A and B compartments, respectively, in both mouse and human cells. They further showed that depletion of L1 RNA reduced A and B compartment strength and their chromatin spatial segregation, suggesting a potentially causative role for L1 in compartment organization. Outside of mammals, however, other repeat classes and families are more prevalent and have been found to correlate with chromatin. In plants such as pepper (*C. annuum*), for example, a strong correlation has been shown between the density of long terminal repeats (LTRs) and A/B compartments and subcompartments [45]. Bredeson et al. found that A and B compartments in the African clawed frog (*Xenopus laevis*) correlated with a linear combination of different repeat element types (specifically the third principal component of repeats), but did not indicate a prominent family, and pointed out the difficulty of determining whether the relationship was causal [24].

The relationship between SINEs, LINEs, and compartments is largely restricted to but also heterogeneous within the mammalian branch (Fig. 2a), as might be expected from the diversity of mammalian SINE families [46]. While primates and rodents showed some of the strongest correlations, Perissodactyla (odd-toed ungulates such as rhinoceros) and Pilosa (e.g., sloths and anteaters) showed reduced or little correlation, in stark contrast to neighboring orders (Cingulata; e.g., armadillos, Fig. 2a). The strong anti-correlation between LTRs and compartments found in the Plantae kingdom is likewise variable in strength, even within the same class (e.g., Magnoliopsida, Fig. 2a). We therefore further investigated the relationship between compartments and repeats by performing a PC analysis of the repeat families (subsets of repeat classes) for each species and compared these principal components with PC1 of the Hi-C contact map, as done by Bredeson et al. [24] for *Xenopus laevis*. In general, the best correlation out of all repeat principal components (linear combinations) with the Hi-C PC1 is comparable to the CDS or gene density correlation with the latter (Extended Data Fig. 4a). However, the interpretability of the repeat PCA is less clear, and a high correlation between the Hi-C PC1 and some linear combination of repeat families, which are in abundance, is not altogether surprising. Regardless, we find for mammals that different phylogenetic orders have implicated different combinations of repeat families with their compartmentalization. Extended Data Fig. 5 summarizes the correlations between repeat families and Hi-C PC1, revealing a lack of a conserved repeat class that shows universal correlation with A/B compartment organization. Instead, different branches have acquired specialized relationships between repeats and their compartment structure.

In addition to the order-level median values shown in Fig. 2a, we show class-level median correlations between these and other genomic features with compartments in Fig. 2b and Extended Data Fig. 4. In the phylogenetic trees in Extended Data Fig. 4b–l, each branch tip is colored by the class-level median correlation value, which we propagated backwards through the tree by estimating maximum likelihood ancestral states with phytools (v2.3.0) and ape (v5.7.1) [47, 48]. While some genomic correlations are unique to certain branches of the tree, such as the mammalian SINE/LINE correlation and the LTR correlation found in plants, others have arisen more than once or in inverted forms. For instance, while a positive correlation with the local GC content is most prevalent in amniotes, a prominent anti-correlation with compartments also arises within many orders of Magnoliopsida, a class of flowering plants (Fig. 2a). To ensure that these correlations are robust, we have determined whether there is any relationship between these correlations and several measures of genome and Hi-C data quality (Extended Data Fig. 6). We find no significant relationship between our main correlations, or their absence, and the data quality. For example, the correlation between GC content and compartments does not depend on the number of experimental Hi-C contacts or genome sequence gap length, even when we focus on non-amniotes. The varied and phylogeny-dependent relationships we have demonstrated suggest a curious history for the use and establishment of compartments throughout the tree of life. Next we use inter-species comparisons to further probe this history.

### Inter-species comparisons to test for similar compartmentalization and non-causality

Fig. 2 reveals the phylogenetically diverse and potentially multi-layered relationship between the genome sequence and its 3D architecture. In mammals, for example, while CDS density is a strong indicator of compartmentalization, local GC content and repeat density can outperform it. It is natural to ask whether these genome sequences correlate strongly because they are playing an active role in the organization of the compartments or have passively aggregated according to an existing structure. Answering such questions experimentally is challenging, since removing large swaths of sequence is not feasible at the scale relevant for compartment organization, and manipulating repeat-associated RNA in a locus-specific manner is not possible with existing approaches. *In silico* methods have been used to probe the effect of removing or adding sequences on compartments [18] and smaller domains [49], but experiments are still required to determine whether the relationships are really causal or simply correlative. While Lu et al. [16] reported that depletion of L1 RNA in mouse embryonic stem cells weakened compartment strength and segregation, compartment identities were largely preserved, leaving the causal role of L1 RNA in compartment formation unresolved.

Here we propose a different test to see which features are non-causal, by using a synteny-anchored comparative analysis. After determining large syntenic blocks shared between two or more species, we simultaneously compare the compartmentalization and the genomic sequence on these shared blocks. If the compartments are similar, which is not obviously guaranteed, but any of the putative predictors are not similar, then it is unlikely that this feature has caused the compartmentalization on these blocks. This may be imagined as an evolutionary knock-in or knock-out experiment, where we let nature reveal the effect of the presence or absence of a specific genomic feature. Stated abstractly, if the raw sequence (*R*) determines an intermediate genomic feature (*F*), and *F* in turn establishes compartment identity (*C*), then whenever *C* is conserved across species, *F* should also remain consistent. Conversely, if *C* is conserved but *F* diverges, *F* cannot be the necessary mediator linking sequence to compartmentalization. This framework thus identifies which genomic features consistently transmit sequence information into stable compartment organization across evolution, and which merely correlate with the Hi-C PC1 in a species-specific manner. While this cannot establish causality, it can rule it out. See Fig. 4a for a schematic of this causal inference in relation to the syntenic block discovery (alignment), and comparison of the genomic features and compartments.

**Fig. 4.**
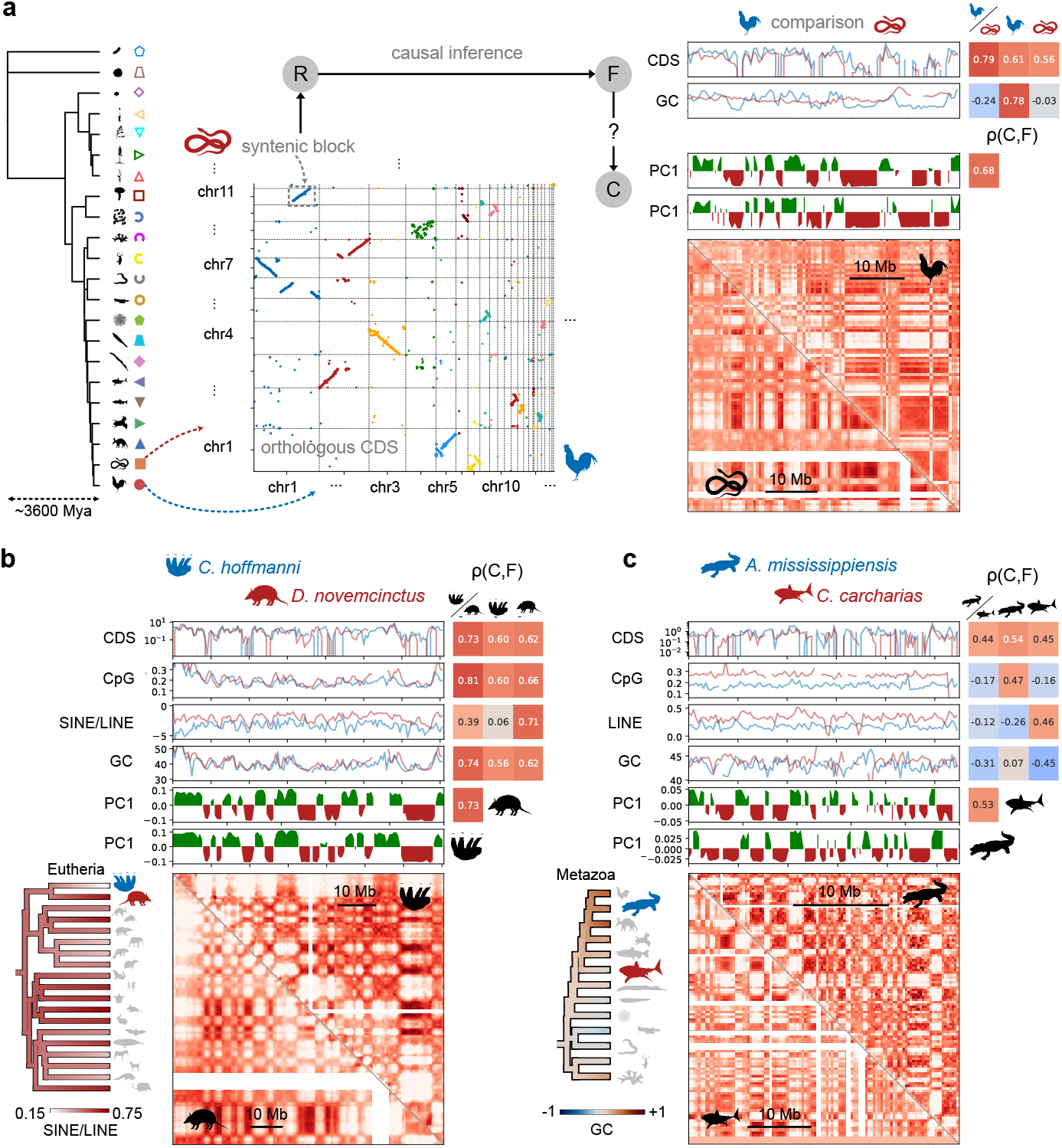
Inter-species comparisons between the Hi-C contact maps, chromatin (A/B) compartments, and candidate genomic features. (a) Schematic of protocol. Two (or more) species are selected from the tree. After aligning similar coding sequences in the two genomes, here shown as a dot plot for multiple chromosomes, large syntenic blocks with similar raw sequence (*R*) are identified for comparison. We then determine whether the genomic features (*F*) and compartmentalization (*C*) are similar on this syntenic block. In the example comparisons, to the right of each genomic feature we include the Pearson correlation of this feature between the species, indicated by the slash, and then the correlation with this feature and the compartments for each species. The upper and lower triangles of the Hi-C contact map belong to the reference and query organisms respectively. Next to each plot we also include a phylogenetic tree with an order-level or class-level ancestral reconstruction [47] of the median Pearson correlation of a particular genomic feature. Here we compared chicken (*G. gallus*) and garter snake (*T. elegans*). (b) Comparison between Hoffman’s sloth (*C. hoffmanni*) and nine-banded armadillo (*D. novemcinctus*). (c) Comparison between alligator (*A. mississippiensis*) and great white shark (*C. carcharias*). In these examples only similarity between CDS consistently predicts similar compartmentalization, indicating that the other genomic features are non-causal innovations.

More concretely, using annotated genomes with corresponding Hi-C contact maps, we used the Oxford Dot Plot (ODP) pipeline [50] with default thresholds to determine conserved orthologous coding sequences shared between species. We then made comparisons of compartment and genomic profiles over syntenic blocks where two or more species share many (≥ 100) orthologous CDSs over a long region (≥ 10 Mb). Relative to a phylogenetic branch with a given consistent compartment correlation, we compare species both within and across this divide to see first whether it is even possible to share similar compartments with different overall correlations. If this prerequisite is fulfilled, we then further probe if this similar compartmentalization required the genomic features to also be similar, at least on these isolated syntenic blocks. This extends previous reports which compared mainly compartmentalization on syntenic blocks [19, 20, 22, 23], by also considering the conservation of multiple genomic features, and by comparing thousands of large syntenic blocks in up to 153 species in and across 7 Vertebrata classes at even greater phylogenetic distances. Here we focus on species that show the typical plaid pattern, as invertebrates and plants showed much higher levels of genome scrambling (see Methods for details regarding synteny block discovery).

### Different compartment rules do not forbid similar compartmentalization

In Figs. 4b–c we show two targeted comparisons of compartments and genomic features that highlight the complexity of conservation. We note the striking conservation of compartmentalization over large (> 50 Mb) blocks between species that are separated phylogenetically not only by hundreds of millions of years, but even by different taxonomic classes, extending previous findings [19, 20, 22, 23] beyond mammals and single classes and to phylogenetic distances over three times larger. Importantly, as we show in Extended Data Figs. 1d–f, such comparisons also demonstrate that compartment sizes are not absolute but scale with chromosome size. In order to also account for inaccurate determination of synteny block coordinates, we thus interpolated one species’s data (assigned the role of query) to the other (reference) species’ data, and allowed for a relative translation (up to 20%) and size scaling (up to 50%), such that this transformation yielded the highest Pearson correlation value with the reference profile. In the targeted comparisons in Fig. 4, we have chosen species residing in different branches of the tree which exhibit different specific correlations with Hi-C PC1 (Figs. 2a, 4a). In these cases, we find striking similarity between compartments, despite different genome-level compartment correlations, and even local, immediate differences on the shared syntenic blocks.

### Inter-species comparisons suggest that genomic features apart from CDS cannot be considered as candidate causal determinants of compartmentalization

For the similar sequence blocks (*R*), where PC1 (*C*) is conserved but individual features (*F*) are not, we can infer that these F are not causally important elements for compartments. In Fig. 4b we show a comparison of a large syntenic block shared between Hoffman’s sloth (*C. hoffmanni*) and the nine-banded armadillo (*D. novemcinctus*). On this syntenic block the Hi-C data is visually quite similar and the compartments are highly conserved (ρ ≃ 0.73). However, while the SINE/LINE density correlates highly with compartments in the nine-banded armadillo (here ρ ≃ 0.71), it is nearly independent in the sloth. Although studies in mice have suggested that LINEs may be involved in the establishment or maintenance of compartments [16], these results suggest that repeats do not play a primary role in causing compartments. This echoes the finding that mobile repeat elements can vary significantly within the similar compartments found in diverse human populations [51].

Other examples from different orders and classes in Fig. 4c and Fig. 5a–c also reveal the phylogenetic diversity of genomic interactions with compartments, but they also demonstrate that genomic features apart from CDS can not generally be considered causal determinants of compartmentalization. The syntenic blocks shown in Fig. 5, which are shared between three species, are particularly pertinent for determining causality. Found by the intersection of syntenic blocks shared between pairs of species, these comparisons on syntenic blocks further highlight how CDS and compartment conservation are strongly tied together independently of the loss or gain of other genomic features that correlate with compartments. In Fig. 5d we move beyond individual examples and perform a general causal inference for CDS, GC, and SINE/LINE density (within mammals) by plotting the Pearson correlation (a measure of conservation) of each feature (*F*) against the correlation (conservation) of the compartments (*C*). This test focuses on the proximity of the data to the diagonal. We supplement the visual comparison with statistical measures included as insets, such as a linear regression slope and the normalized root mean square deviation. Only the conservation of CDS consistently indicates the conservation of Hi-C PC1, indicating that GC and SINE/LINE density (in mammals) are not causative rules for compartmentalization but are most likely themselves a product of the gene density and corresponding compartmentalization. Additional evidence that the conservation of CDS controls compartmentalization is provided by restricting attention to orthologous (conserved) CDS. For each syntenic block in Fig. 5d we also calculated the density of the CDS that are shared between the two species, according to the default thresholds used in the ODP pipeline [50]. When compared with the compartments on these same blocks, the density of conserved CDS significantly outperforms the full CDS density (Fig. 5e). This further links nuclear compartmentalization with an inherited gene density.

**Fig. 5.**
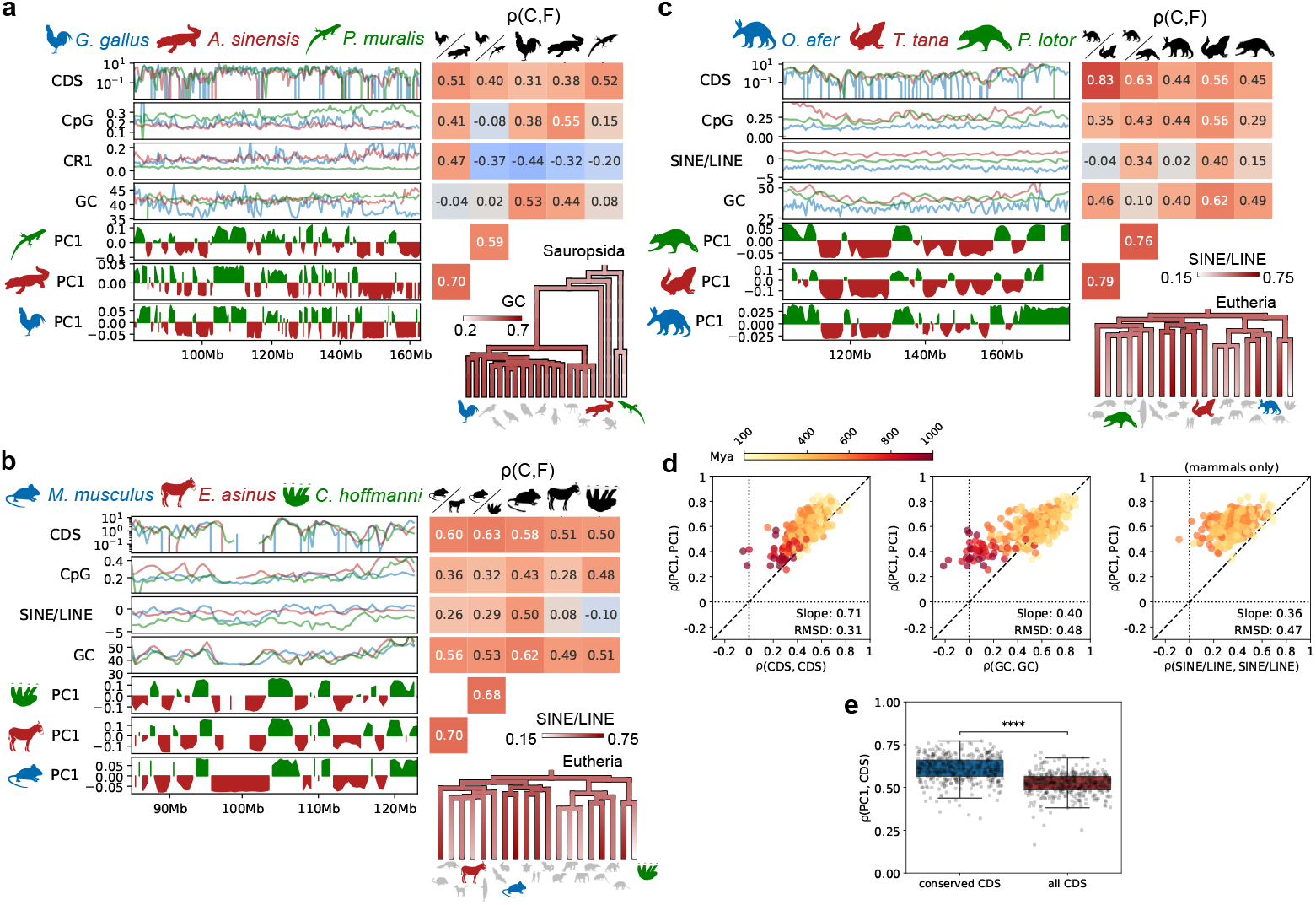
Three-species comparisons of genomic features and compartmentalization on large syntenic blocks and a global test of causality. As in Fig. 4b–c, we show Pearson correlation between the two query species’ and one reference species’ data to the right of the data plots in the first two columns, and the correlation between this data track and the Hi-C PC1 for each species separately in the last three columns.(a) Comparison of chicken (*G. gallus*), Chinese alligator (*A. sinensis*), and wall lizard (*P. muralis*). (b) Comparison of mouse (*M. musculus*), donkey (*E. asinus*), and Hoffmann’s sloth (*C. hoffmanni*). (c) Comparison of aardvark (*O. afer*), large tree shrew (*T. tana*), and raccoon (*P. lotor*). These targeted comparisons show that where CDS and PC1 are conserved, the GC content and SINE/LINE density can be plastic. (d) Inter-order median plots within vertebrates of the Pearson correlation of different features and PC1 on syntenic blocks (shared by two species). Consistent prediction is measured by the data’s nearness to the diagonal dashed line. Of the three candidates, CDS density predicts conservation of PC1 with most consistency. Within each subplot we include statistical diagnostics of the ability of each feature to predict the Hi-C PC1, such as a linear regression slope and the normalized root mean square deviation. (e) A comparison of the correlation between Hi-C PC1 and the conserved CDS density vs. the full CDS density on the syntenic blocks from (d). Restricting attention to conserved CDS significantly increases the correlation with compartmentalization.

## Discussion

In this study, we discovered that the A/B chromatin compartments elucidated by a principal component analysis of Hi-C contact maps are most intimately and generally connected with CDS density. The distribution of these sequences explains the genome folding at the sub-chromosomal scale not only in organisms with the familiar plaid-type Hi-C contact maps but also in species with non-plaid, monochromatic Hi-C maps. Using inter-species comparisons to rule out non-causal factors, we find that the CDS density landscape remains the only robust candidate among the usual suspects proposed to drive chromatin compartmentalization. Different branches of the evolutionary tree may have thus used their inherited gene density landscape and associated genome folding as a scaffolding upon which new and diverse relationships between genome sequence features and 3D chromosome organization were subsequently built. This may be regarded as a large-scale version of the well-known notion that gene regulatory sequences evolve more quickly than the genes they regulate [52].

A central question raised by these findings is what mechanistic link connects CDS density to spatial compartmentalization. One plausible candidate is transcription itself. Active genes recruit RNA polymerase II and associated factors, and the clustering of co-transcribed loci into shared transcription hubs has long been proposed to drive spatial co-segregation of active chromatin [53, 54]. Consistent with this view, A compartments are tightly associated with nuclear speckles—membraneless condensates enriched in RNA-processing factors—which broadly co-localize with gene-dense, transcriptionally active regions across cell types and species [55, 56]. Developmental dynamics further support a potential role for transcription in establishing large-scale compartment architecture. In early mouse embryos, the emergence of a somatic-like replication timing (RT) program occurs shortly after zygotic genome activation (ZGA), when widespread transcription is first initiated after fertilization [57]. Because replication timing strongly correlates with A/B compartments, the near-synchronous establishment of genome-wide transcription and somatic-like RT suggests that compartmentalization may likewise emerge in close association with global transcriptional activation. From this perspective, transcription could serve as a functional bridge linking the static genomic landscape of CDS density to dynamic nuclear compartment organization.

Independently, CDS-rich regions are preferentially marked by histone acetylation (H3K27ac, H3K9ac), which directly loosens chromatin compaction [58, 59], and is among the most broadly conserved epigenomic features across metazoans [52]. The precise molecular details likely vary across the tree of life, however, as nuclear speckles are not universal and histone modification landscapes can differ substantially between clades [60, 61]. Nonetheless, these mechanisms collectively suggest that CDS density, by directing the formation of transcriptional hubs and establishing a self-reinforcing chromatin modification environment, can act as a primary determinant of compartment identity, with other genomic features arising secondarily as a consequence.

A possible mechanism for heterogenizing genomic features at the level of compartment domains that relates to both the enhancement of GC content in amniote compartments, the repeat landscape in mammalian compartments, and the plaid-monochromatic Hi-C contact patterns, is genetic recombination [62]. Occasional genetic recombination reshuffles the genetic material on the chromosomes in offspring sometimes by the physical breaking and rejoining of parent DNA strands. Recombination does not occur uniformly in chromosomes, but tends to concentrate in regions with a higher density of genes [63] and open chromatin [64, 65], and organisms tend to retain healthy gene mutations while eliminating deleterious ones [66]. High recombination rates are known to be concentrated at the ends of chromosomes in many plants [67], while they are heterogeneously distributed in mammalian genomes [64], thus reflecting the plaid-monochromatic behavior seen in Hi-C contact maps and CDS density. Within mammals and birds, there is an error correction bias (i.e., GC-biased gene conversion or gBGC), which can tend to increase GC content especially in regions with high recombination rates [68]. Thus the inherited chromatin compartmentalization, combined with this correction bias, may have led to the strong GC content correlation broadly observed in amniotes. Further investigation into the relationship between recombination, genome features such as repeat elements [69], and genome architecture should shed further light on these issues.

These findings may help explain the phylogenetic diversity observed in chromosome banding patterns [1, 2]. The general consensus, at least for earlier techniques, was that intricate G-banding (or R- or Q-banding) is largely restricted to birds, mammals, and possibly reptiles. This is presumably due to the presence of isochores, the strong heterogeneity of GC content which varies along the length of each chromosome. These isochores are ubiquitous in mammals and birds but rare in most cold-blooded vertebrates [3, 70]. We found that the correlation between GC content and chromatin compartments is largely restricted to amniotes (see Figs. 2a, 3d), suggesting that the compartment structure induced by CDS density variations created isochores in this group, yielding distinct G-bands, but not in other clades. Using the same length-scale analysis employed in Fig. 3, we likewise found a significant difference between the ratio of characteristic length scales for GC and Hi-C PC1 within and outside of amniotes (see Extended Data Fig. 7). In amniotes the GC structure (isochores) appears to follow but not cause compartmentalization (see Figs. 4, 5). In those species where the heterogeneity is sufficient, chromosome banding patterns may also be observed, but these bands are not the most faithful indicator of physical compartmentalization.

Our work ultimately redirects attention to an upstream determinant of nuclear architecture, namely CDS density heterogeneity along chromosomes. Across species with a wide range of genome sizes, we identify two extreme and distinct genome architectural styles, plaid and monochromatic. Our phylogenetic analysis suggests that monochromatic genome architecture has arisen independently multiple times, implying that it represents a recurrent evolutionary outcome. Our results further suggest that these architectural styles reflect the genomic distribution of coding sequences shaped by lineage-specific evolutionary histories. The architectural style of physical buildings, such as baroque or minimalist, is likewise a product of history and geography, yet all buildings remain constrained by universal mechanical principles. In the same way, variation in genome architectural style may emerge from lineage-specific patterns of CDS distribution operating within shared physical constraints on chromatin organization. From this perspective, plaid compartment organization appears not as a fundamental organizing principle, but as an emergent structural outcome of genome evolution.

The substantial diversity of Mb-scale chromatin organization across species further suggests that nuclear compartmentalization is subject to relatively weak evolutionary constraint. This is consistent with the observation that gene regulatory interactions, such as promoter-enhancer contacts, occur at much smaller genomic scales. Interestingly, these local chromatin contacts have been shown to remain largely intact despite variation in Mb-scale genome folding, for instance, by condensin II depletion in human cells [6]. From this perspective, Mb-scale compartment architecture may be free to diverge as long as local regulatory logic is preserved, allowing lineage-specific CDS density landscapes to be directly reflected in lineage-specific nuclear organization.

While the relationship between small-scale TADs and syntenic breakpoints has been investigated numerous times [21], previous work has also found that syntenic breakpoints can correlate with chromatin compartments [71]. However, these investigations have remained limited in scope. We speculate that many of these findings may be modified when similar analyses are conducted across other branches of the tree of life. Here we have demonstrated the power of a broad phylogenetic comparison to eliminate many of the usual genomic suspects, highlight important differences, and (re-)focus our attention on the upstream cause, the gene density heterogeneity along each chromosome.

## Methods

### Preparing and analyzing the genomes

We collected chromosome-level genomes and corresponding Hi-C data mainly from NCBI [25], DNA Zoo [26, 27], and GenomeArk [28]. Genomes hosted on DNA Zoo often have Hi-C contact map data already aligned to the genome assembly. In all other cases we aligned the raw Hi-C reads to the genome assemblies using the Juicer platform [27] with default parameters. We show that our main results, namely the correlation between CDS density and Hi-C PC1, do not depend on the quality of the Hi-C data in Extended Data Fig. 6. In the case of the large tree shrew (*Tupaia tana*) belonging to the Mammalian order Scandentia we used the scaffold-level assembly (GCA 004365275.1) and the Dovetail Hi-C reads (BioProject PRJNA782001) available on the NCBI database and then assembled the genome *de novo* using the 3D-DNA pipeline [27]. If a genome was not annotated, or if its existing annotations’ BUSCO completeness score was low, we generated annotations using GeMoMa (v1.9) [30], Liftoff (v1.6.3)[31], or both, with high quality annotated genomes of closely related species. Multiple annotations were merged using AGAT (v0.8.0) [72] and typically achieved annotation BUSCO completeness scores higher than 90% as determined by BUSCO (v5.2.2) [29]. The local GC content was determined using Biopython (v1.79) [73], and the CpG density was determined using the method of Gardiner-Garden et al. [74]. We searched for repeat elements using RepeatMasker (v4.1.8) with the rmblast engine (a RepeatMasker-optimized version of BLAST+) [33] after determining *de novo* repeat family libraries with RepeatModeler (v2.0.6) [32]. More details on finding repeat elements can be found below. We excluded individual chromosomes that had apparent assembly errors or other defects that affected the Hi-C data. Small sex chromosomes such as chrY in mouse, for example, were typically excluded. See database and Extended Data Fig. 8. Statistical comparisons were made using scipy (v1.5.3) [75], and the statistical annotations shown in Fig. 2b, Fig. 5e, and Extended Data Fig. 7a were created using statannotations (v0.6.0) [76].

### Identifying plaid and monochromatic chromosomes

We determined whether a chromosome was plaid or monochromatic based on the ratio of its Hi-C PC1 correlation length to the chromosome size, which was typically less than 0.2 for all chromosomes. A probability distribution of this ratio is shown in Ext. Data Fig. 2a for all species and all chromosomes. The animal kingdom, which contains most of the data in our database, is largely confined to smaller ratios. The non-animal kingdoms’ length scales are more dispersed and extend to higher fractions of the chromosome size. We chose a fraction of 0.12 as a threshold to identify plaid and monochromatic chromosomes. We then defined plaid/-monochromatic species as those with more than or equal to 2/3 of their chromosomes having the respective plaid/monochromatic character. Species which satisfied neither of these criterion were mixed. We then determined whether a taxonomic order was plaid/monochromatic based on the same 2/3 criterion for its species in our database.

### Assembling the large tree shrew (*Tupaia tana*) genome

In order to maximize the mammalian orders represented in our analysis we improved the assembly of the current scaffold-level genome assembly of *Tupaia tana* (GCA 026018925.1), belonging to the order Scandentia, using Dovetail Hi-C reads (PRJNA782001) [71]. We aligned the raw Hi-C reads to the genome assembly using the Juicer pipeline (v1.6), made a new draft assembly from the scaffolds with the 3D-DNA pipeline (run-asm-pipeline.sh, version 180419), manually curated the scaffolding using the Juicebox Assembly Tools, and then polished them again with the 3D-DNA pipeline (run-asm-pipeline-post-review.sh) [27]. For our final analysis we excluded all scaffolds smaller than 10Mb, leaving us with a genome of 29 chromosomes amounting to ∼ 2.68Gb, which is comparable to species in nearby orders (*Mus musculus*, 21 chromosomes, ∼ 2.75Gb). We then re-ran the Juicer pipeline on the chromosome-level assembly to obtain Hi-C contact maps and annotated the genome as described above. The final genome assembly annotations achieved a BUSCO completeness score of 95.1% (mammalia odb10).

### Synteny-anchored comparisons

We aligned coding sequences using the ODP (Oxford Dot Plot) pipeline [50], which filters and combines blastp (v2.15.0) [77] results to produce a reciprocal best hit shared list of orthologous proteins along with their corresponding CDS genomic coordinates. While originally developed to determine shared (ancient) syntenic regions from multiple species, here we use it for the much simpler task of identifying orthologous CDS and syntenic regions between two species. We used the pipeline’s default thresholds and further filtered and extracted syntenic blocks using custom Python scripts. The syntenic blocks were constructed using coding sequence conservation and collinearity rather than overall sequence composition. Because this pipeline relies specifically on coding sequence, it isolates inherited CDS density patterns independently of lineage-specific variation in GC content or repeat landscapes. We additionally used LASTZ [78], following the same procedure as Corbo et al. [23], to align genomes to confirm the results in a limited number of cases (See Extended Data Fig. 9). In these comparisons one species is assigned the role of reference and the other as query. In all cases the query species’ data was interpolated onto the reference species’ genomic coordinates. Because we do not expect these coordinates to be exact and because of relative genome size variations, we allowed for up to 20% relative translations and scaling up to 50% of the ratio of the genome sizes and chose the transformation of the query compartment profile which yields the highest Pearson correlation value with the reference profile. In our final analyses shown in Fig. 4 and Fig. 5 we restricted attention to species in Vertebrata, and considered only syntenic block pairs for which ρ(*C, C*) is positive and the p-value is less than 0.05. The statistical measures shown as insets in Fig. 5d were computed using an even phylogenetic re-sampling procedure to avoid over-representation of closely related species, which contributed disproportionately more comparisons and syntenic blocks. Pairwise comparisons were first grouped into bins by phylogenetic distance, and an equal number of samples was randomly drawn from each bin. This procedure was repeated 100 times, and the reported values represent the mean across all re-sampled datasets.

### Phylogenetic tree

Phylogenetic distances between species were determined with TimeTree (v5) [34]. When species were not found in this database, we took a sister taxa which was not more closely related to the other species in our search. For *Strix occidentalis* (spotted owl) [79] and *Halictus rubicundus* (orange-legged furrow bee) [80] we manually added a branch using phylogenetic distances found in the literature. We used MEGA (v11) [81] and FigTree (v1.4.5) [82] to manipulate and plot the phylogenetic trees in Figs. 1, 4.

### RepeatModeler and RepeatMasker

We performed *de novo* searches for repeat elements using RepeatModeler (v2.0.6) [32] with the optional LTR strucural finder (-LTRStruct flag) for all species. The search was performed only on the largest chromosome except when memory constraints required a smaller chromosome. The *de novo* library was then used to determine repeats throughout each species’ genome using RepeatMasker (v4.1.8) [33]. We also determined repeats using the existing libraries (Dfam with RepeatMasker Reference Models) and compared against our *de novo* results. As shown in Extended Data Fig. 10, in general the two libraries produced similar findings, but significantly more LTR elements were found using the *de novo* libraries especially in plants, resulting in generally higher correlation with the Hi-C PC1.

### Silhouette and Images

The silhouette images used in the figures were all obtained from www.phylopic.org. The silhouette of *Dromiciops gliroides* used for Microbiotheria and the silhouette of *Notoryctes typhlops* used for Notoryctemorphia were created by Sarah Werning. Unless otherwise noted, all other images are in the public domain.

### Data, Materials, and Software Availability

The tables of genome and Hi-C data sources, analysis parameters, correlation values, analysis code and other relevant information supporting the findings of this study has been deposited in the Zenodo open repository at https://doi.org/10.5281/zenodo.18332941 under the MIT License and is available to reviewers and editors during peer review. Access will be made fully public upon publication. All genomic and Hi-C data were obtained from public repositories.

## Acknowledgments

We are grateful to all members of the Hiratani and Kawaguchi laboratories for their insightful discussions and feedback throughout this project. In particular, we thank Jothivanan Elumalai, Linda Choubani, and Yohsuke Fukai for their careful reading of the manuscript. We acknowledge support by the RIKEN Information systems division for the use of the Supercomputer HOKUSAI BigWaterfall. We also acknowledge the use of large language models, specifically ChatGPT (OpenAI), to assist with the editing of the manuscript text, as well as for troubleshooting code during data analysis. The authors reviewed, edited, and take full responsibility for all content.

## Author contributions

R.T.C., K.K., and I.H. conceived the study. R.T.C. designed and performed the analysis. The manuscript was written by all co-authors.

---

**Extended Data Fig. 1.**
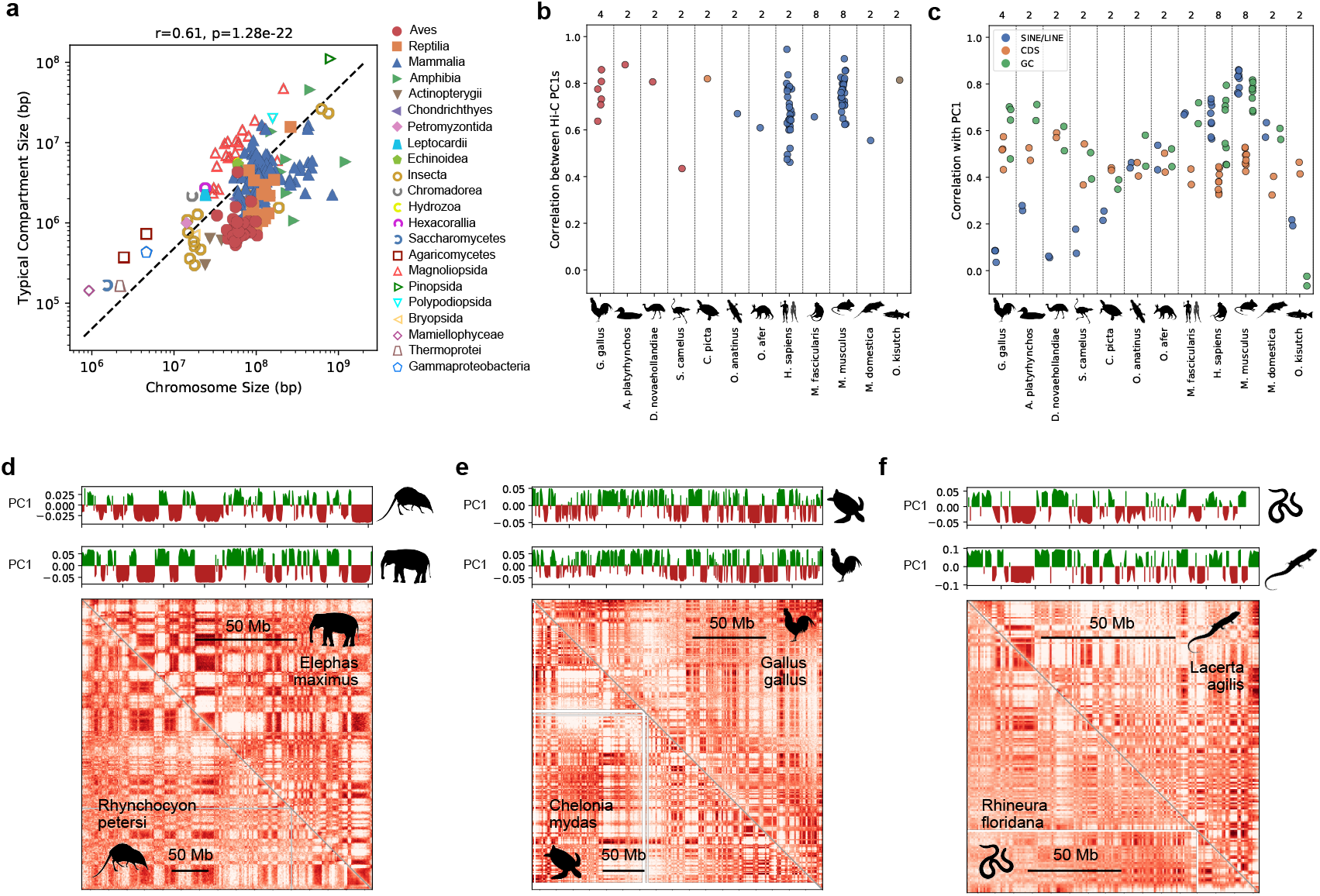
Scaling of compartments and TADs with genome size. (a) Median typical compartment size plotted versus median chromosome size. The compartment length scales were extracted using the autocorrelation of the A/B compartment (PC1) profile for each chromosome in a species. The length scale was determined as the location when the autocorrelation fell below a third of its maximum value. (b) Correlation between Hi-C PC1 values within a species for different cell types (tissue samples). The number of cell types for each species are denoted above. (c) Median Pearson correlation between Hi-C PC1 and CDS density, local GC content, and SINE/LINE density for species with multiple cell type data. Notwithstanding the variation, the overall trends are consistent with the findings presented in Fig. 2a. The number of cell types for each species are denoted above. (d)-(f) Examples of similar Hi-C data and A/B compartments on syntenic blocks between species with different size genomes. Even when the relative sequence size is doubled, and the tissue samples (cell types) are different, the compartments can show remarkable similarity, highlighting the chromosome size dependence of the compartment sizes.

**Extended Data Fig. 2.**
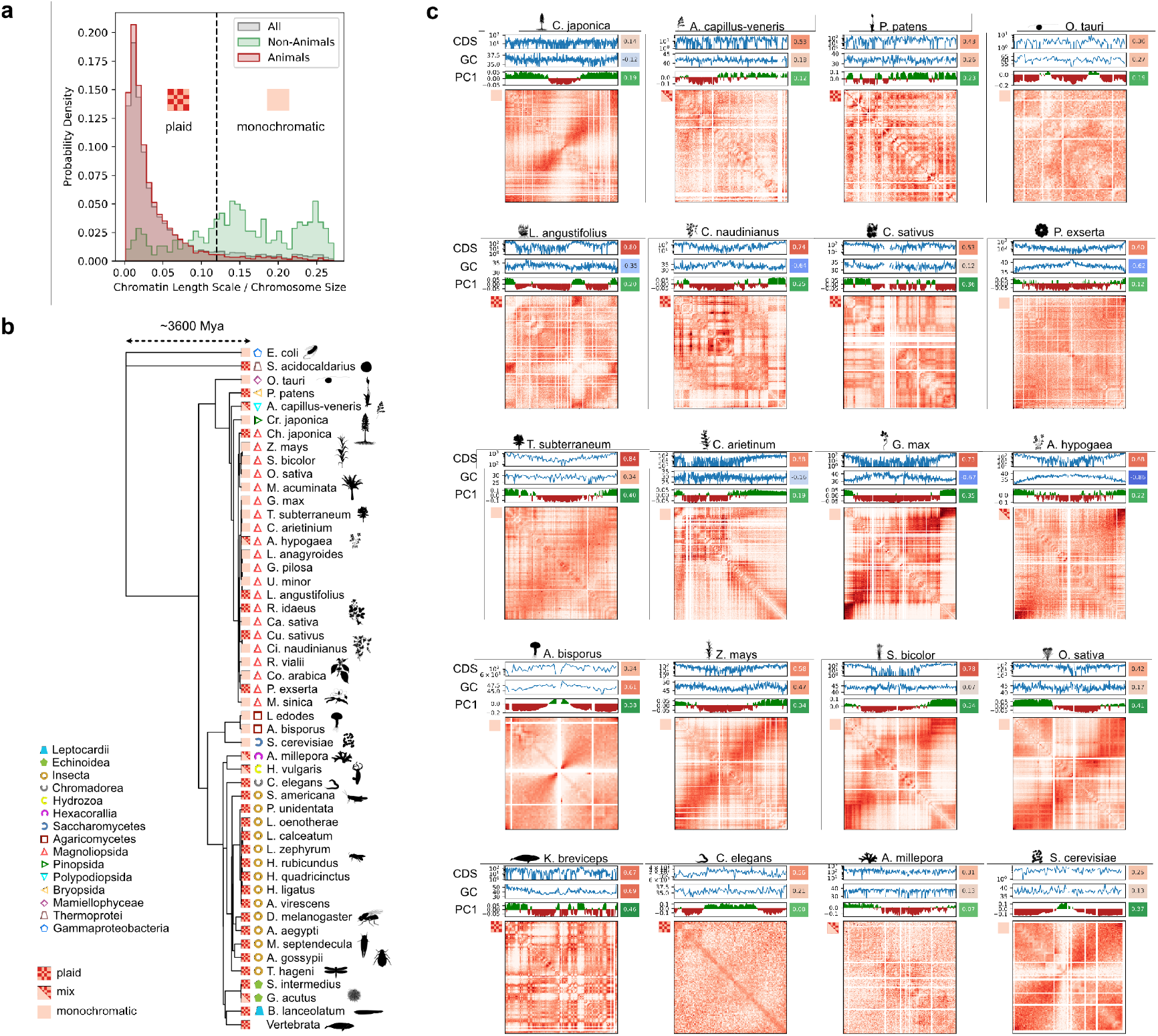
More details on the phylogeny and diversity of plaid and monochromatic genome architectures. Nearly all vertebrates studied showed a plaid Hi-C contact pattern. (a) Probability distribution of the chromatin (correlation) length scale normalized by the chromosome size for all species and all chromosomes. The animal kingdom is closely clustered at very small scales. The non-animal kingdoms’ length scales are more dispersed and extend to higher fractions of the chromosome size. We chose a fraction of 0.12 as a threshold to identify plaid and monochromatic chromosomes and then defined plaid/monochromatic species as those with more than or equal to 2/3 of their chromosomes having the respective plaid/monochromatic character. Species which satisfied neither of these criterion were mixed. (b) Phylogeny of all species with a magnification of the non-vertebrate portion of the tree revealing the phylogenetic complexity of the plaid-monochromatic trait. (c) Examples of all monochromatic species as well as several interspersed plaid and mixed species. While the CDS density consistently correlates even in monochromatic species, the local GC content can either hardly correlate with Hi-C PC1 or sometimes even show the opposite tendency as the CDS density.

**Extended Data Fig. 3.**
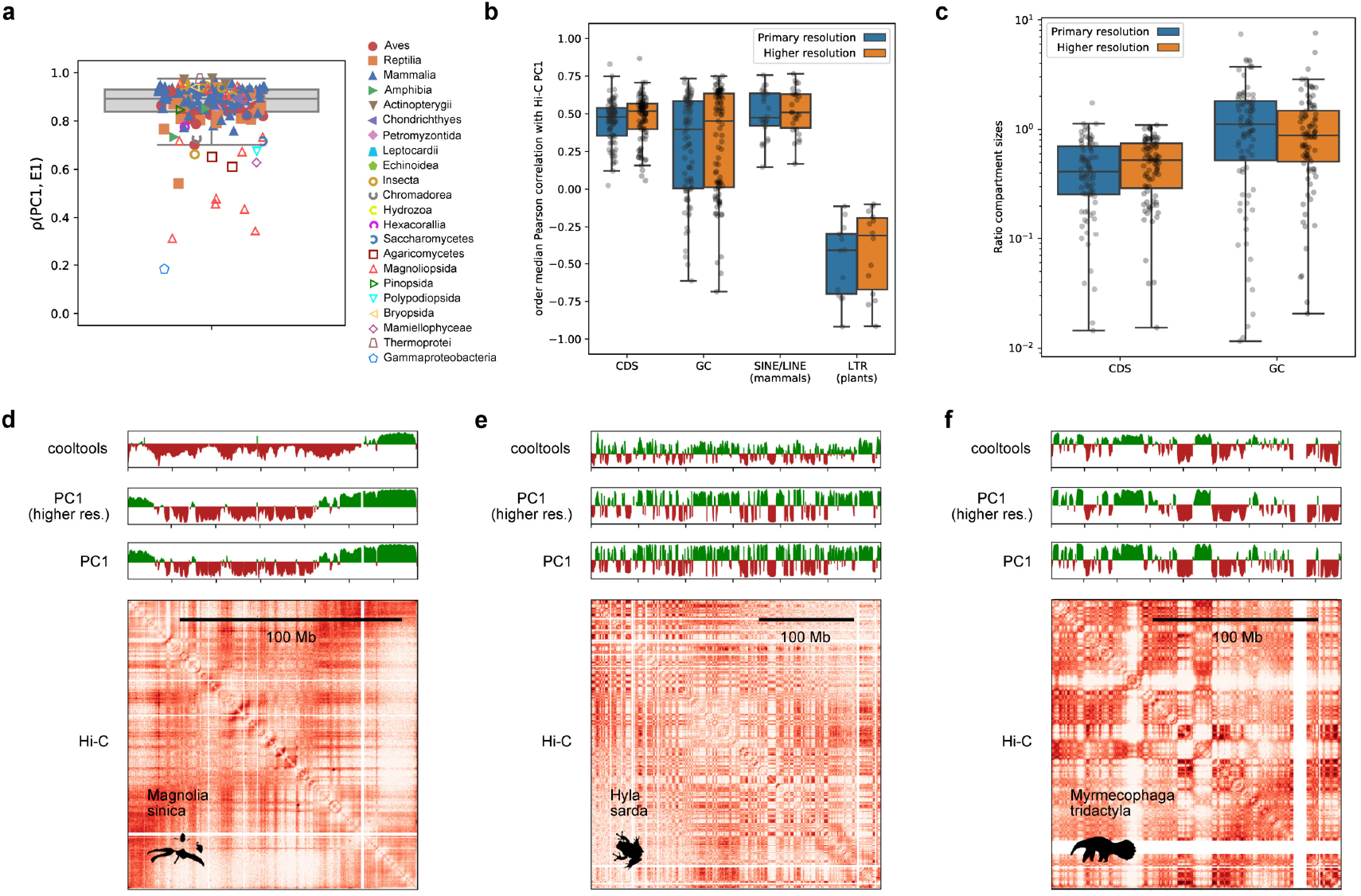
Tests of the robustness of the compartment calling method. (a) Comparison between the current (similar to Lieberman et al. [4]) method of estimating compartments (PC1) and the result from the *cooltools* package (E1). (b) Comparison between the correlation estimates for the resolution used in the primary analysis and the next highest available Hi-C resolution. (c) Comparison between the ratio of autocorrelation length scales (CDS and GC to Hi-C PC1) for the resolution used in the primary analysis and the next highest available Hi-C resolution. (d)-(f) Hi-C contact map and compartment profiles estimated using the resolution in the primary analysis (PC1), the next highest available resolution (PC1, higher res.), and cooltools at the resolution of the primary analysis (cooltools) for (d) huagaimu (*M. sinica*), leaf sample, chr4; (e) sardinian treefrog (*H. sarda*), muscle sample, chr6; and (f) giant anteater (*M. tridactyla*), blood sample, HiC_scaffold_2.

**Extended Data Fig. 4.**
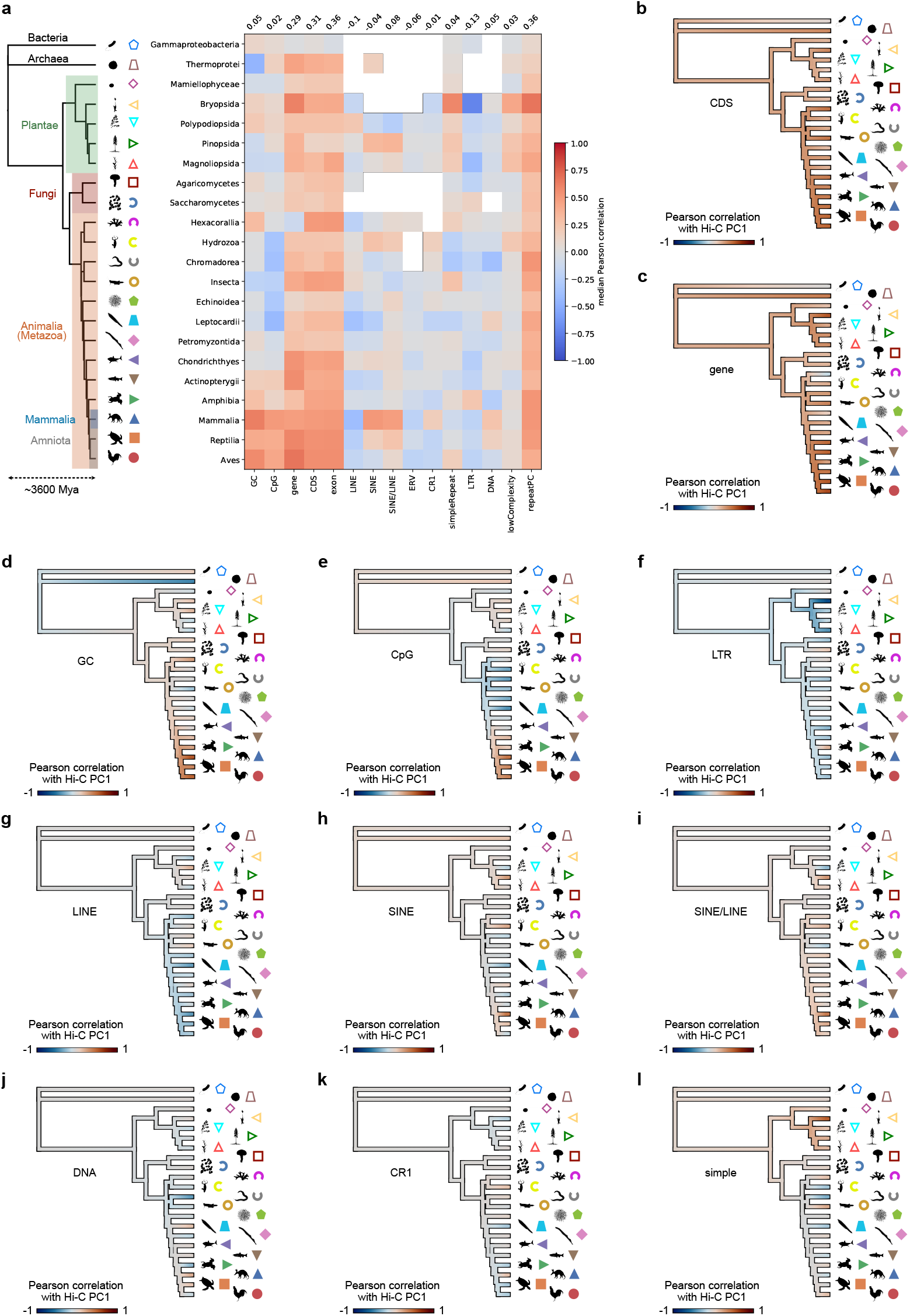
Summary of correlations between Hi-C contact map’s PC1 and various genomic features. (a) A heatmap of the median Pearson correlation at the class level and arranged according to the phylogenetic tree. Overlaid over the tree are indications of certain clades. (b)-(l) Phylogenetic trees at the class level showing estimates of maximum likelihood ancestral states for the median correlation between the Hi-C PC1 and various genomic features.

**Extended Data Fig. 5.**
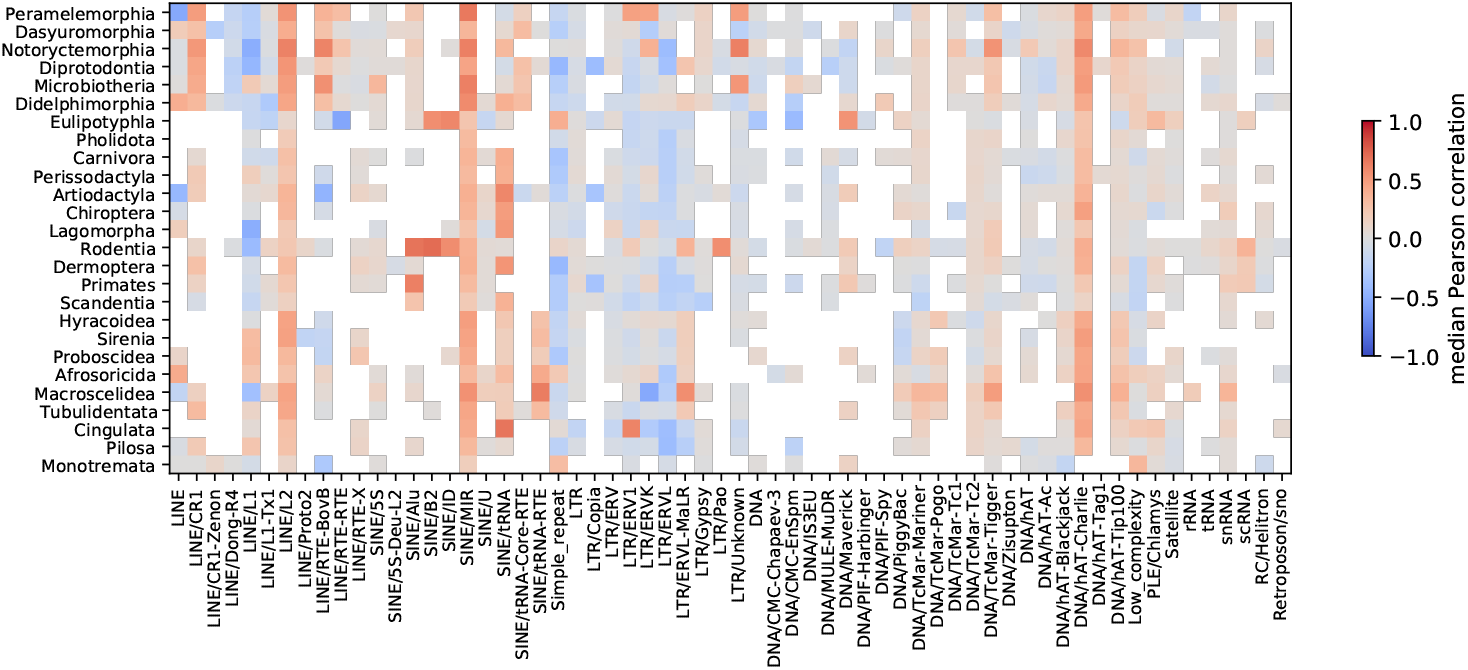
Summary of correlation between repeat classes and families with PC1 for Mammalia. The diversity observed in the correlation between PC1 and the ratio of all SINEs over all LINEs shown in Fig. 2 is a reflection of the diversity of correlations among different repeat families. Moreover, in orders where SINEs and LINEs correlate weakly with PC1, other repeat types such as DNA repeats emerge as a dominant contributor.

**Extended Data Fig. 6.**
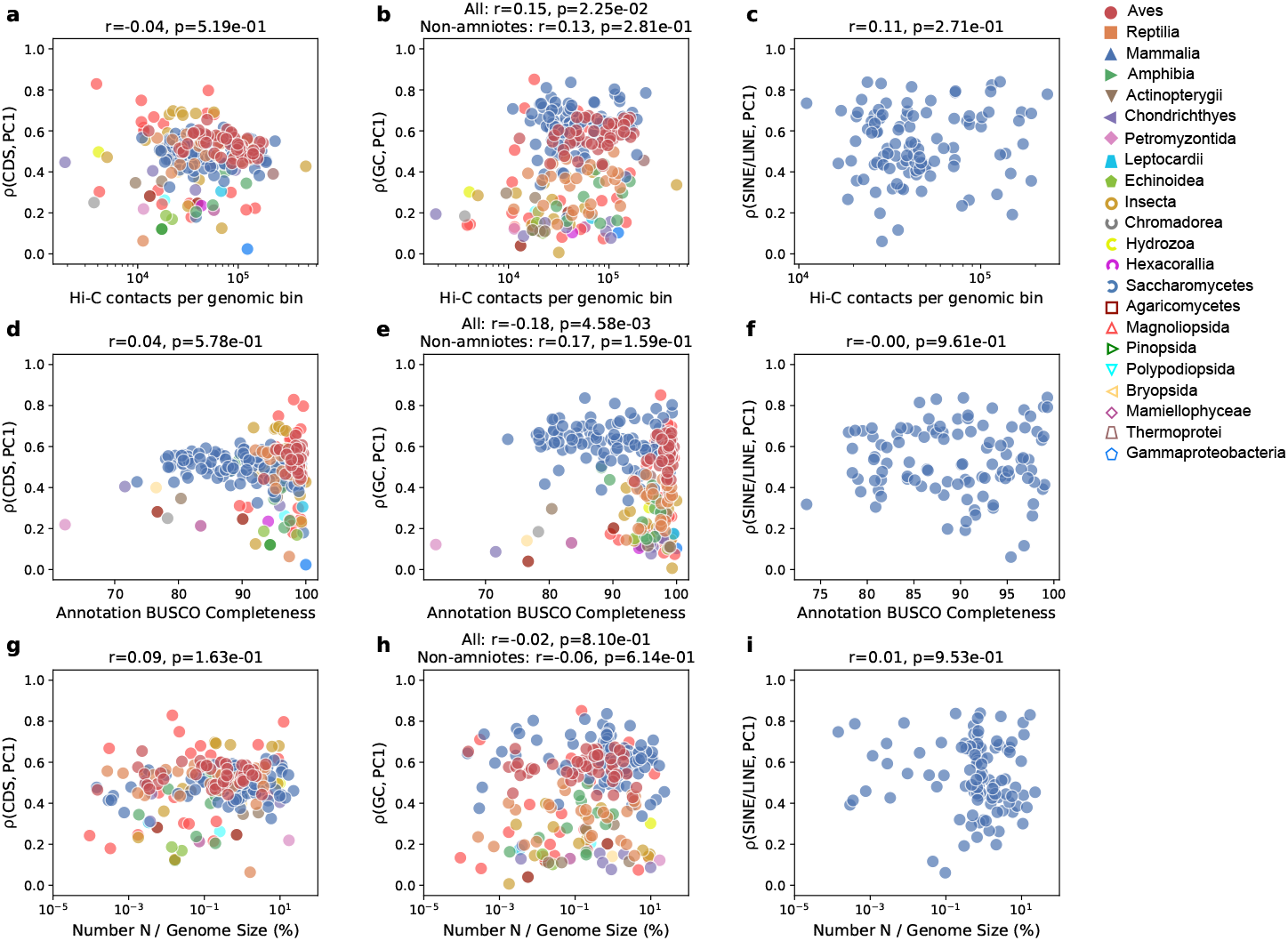
Testing dependence of correlations with CDS, GC, and SINE/LINE (in mammals) on genome and Hi-C data quality. All correlations are given for species median values.

**Extended Data Fig. 7.**
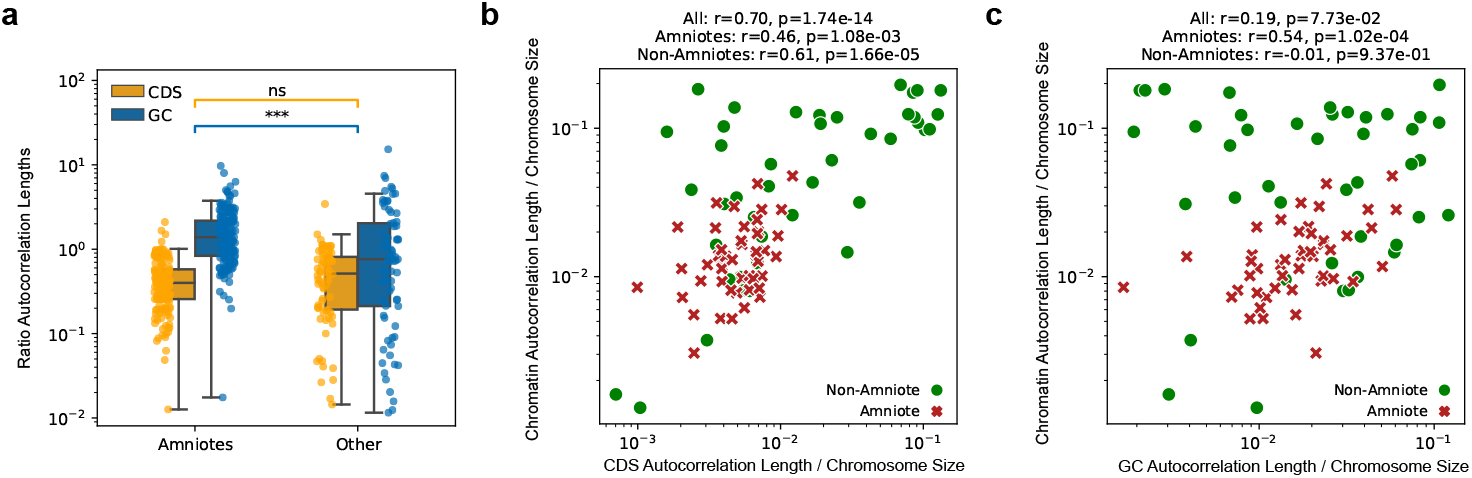
(a) Plot of ratio of CDS and GC typical compartment size to the typical chromatin compartment size distinguishing between amniotes (mammals, birds, and reptiles) and non-amniotes. Chromosome (G-)banding is known to be more prominent in amniotes, and we likewise find that amniotes have a tighter and statistically different relationship between chromatin compartments and the GC content compartments which may give rise to mitotic banding patterns. (b)-(c) Species-level median plots of chromatin compartments vs. CDS and GC content typical compartment sizes. These results follow the same trend as the order-level plots in Fig. 3.

**Extended Data Fig. 8.**
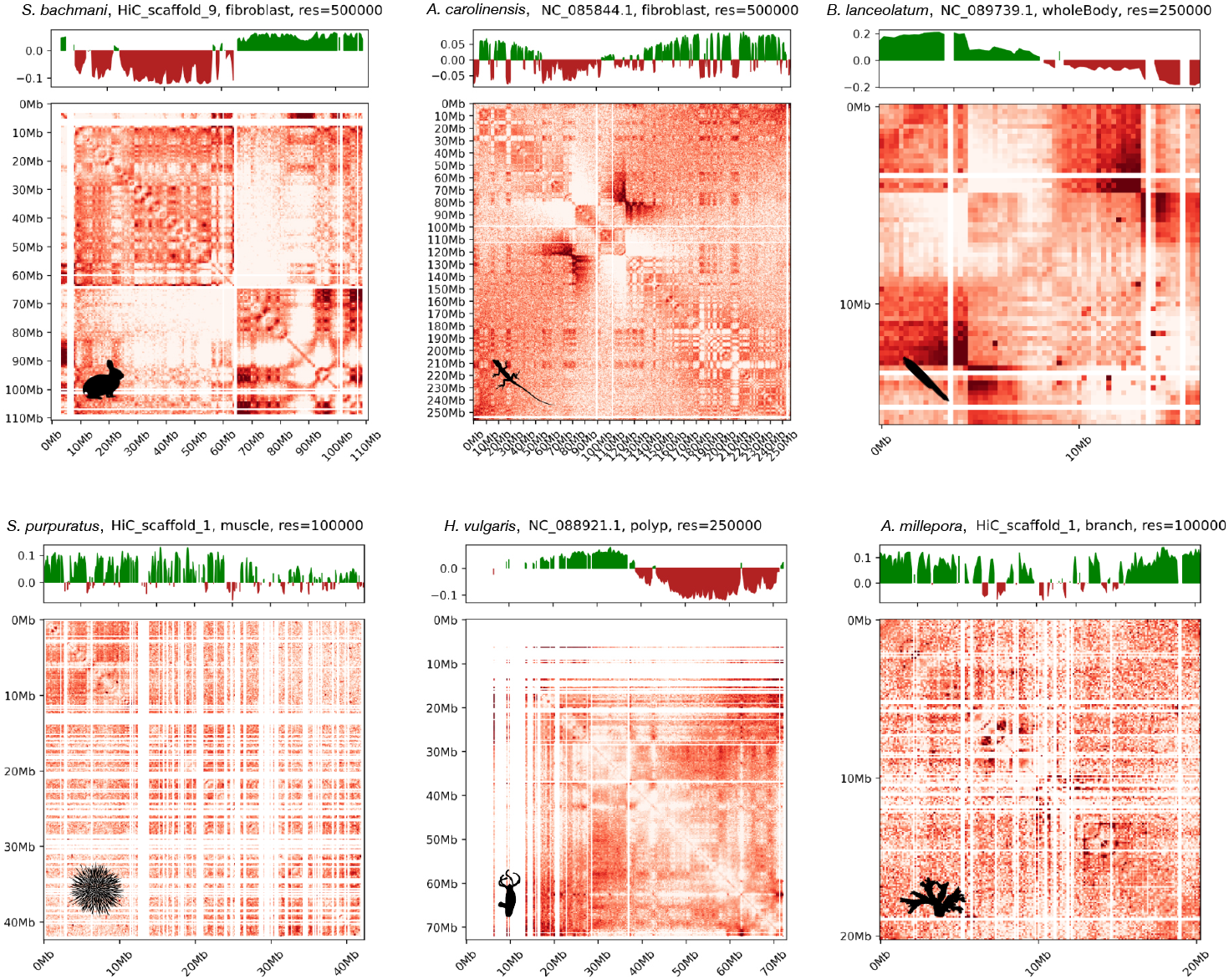
Examples of individual chromosomes which show abnormal contact patterns, apparent assembly errors, or sufficiently extensive unmapped regions such that the character of PC1 is adversely affected.These chromosomes are identified manually and excluded from the analysis.

**Extended Data Fig. 9.**
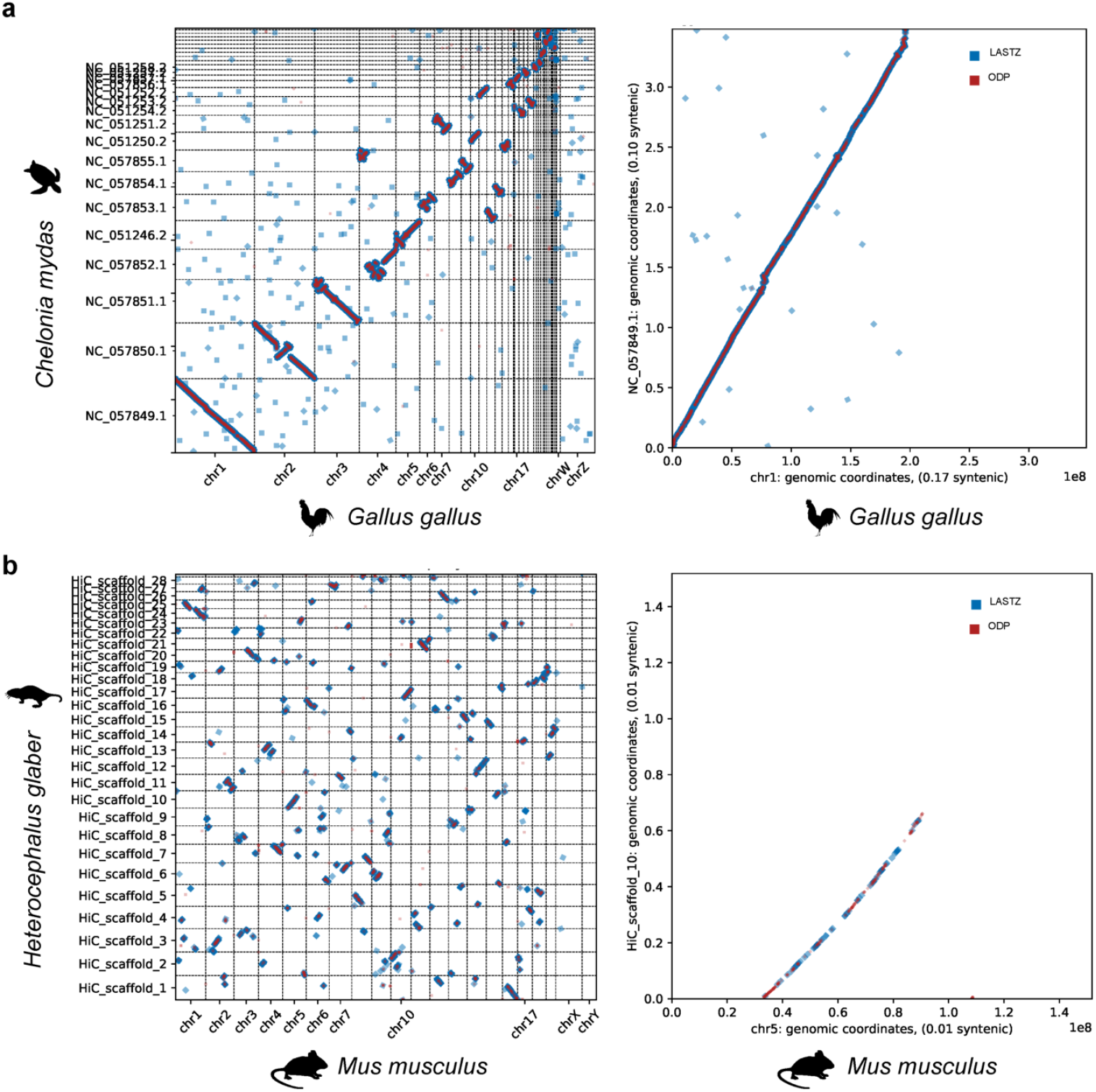
Confirmation that the ODP pipeline [50] yields syntenic regions similar to that of a LASTZ pipeline [23, 78]. (a) Chicken (*G. gallus*) vs. sea turtle (*C. mydas*). (b) Mouse (*M. musculus*) vs. naked mole rat (*H. glaber*).

**Extended Data Fig. 10.**
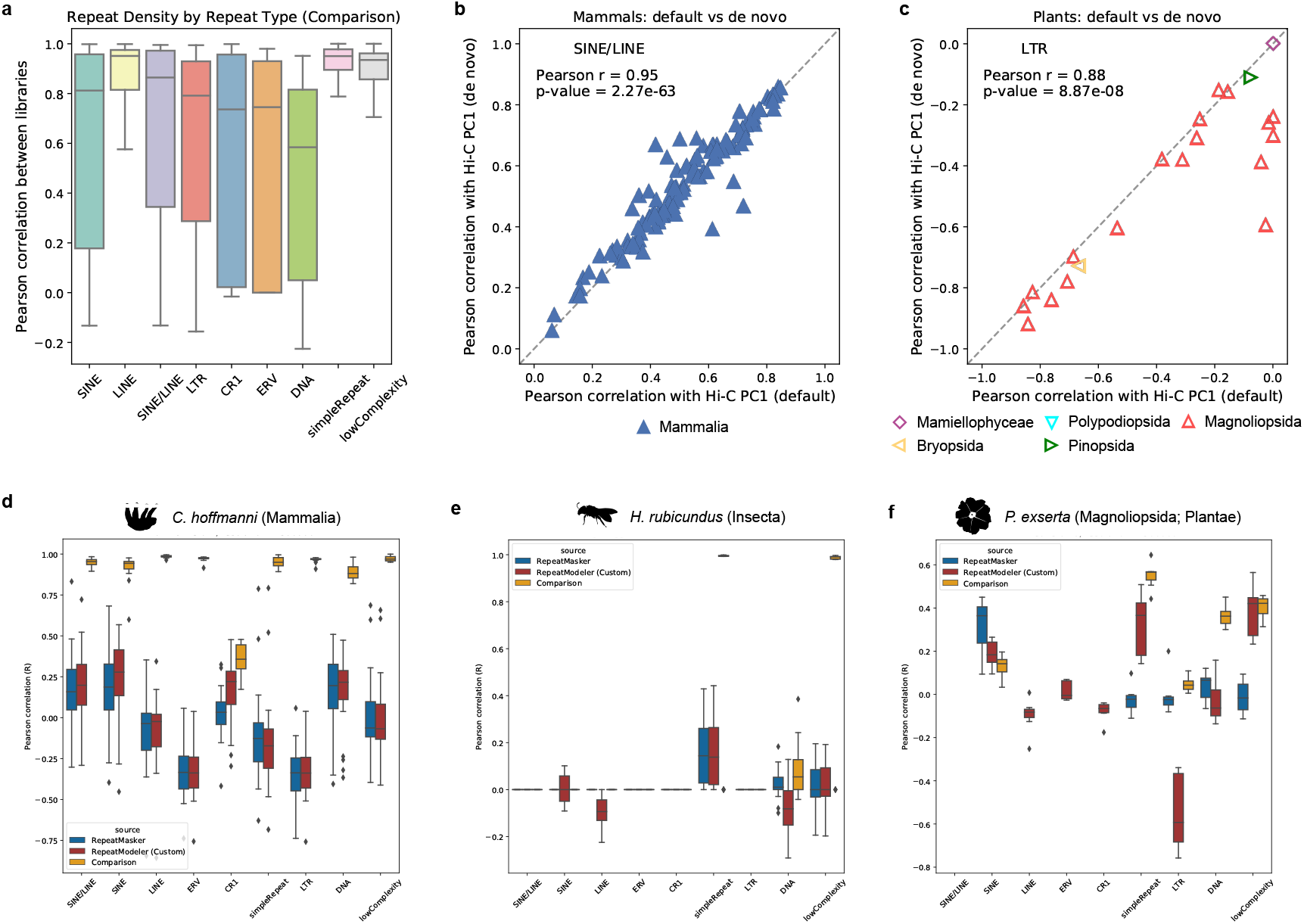
Comparison of estimates of repeat density. (a) Box and whisker plot summary of Pearson correlations between repeat densities within a species using existing (default) RepeatMasker libraries or custom (*de novo*) libraries generated with RepeatModeler. We also plot the Pearson correlation of each with the Hi-C PC1 for the SINE/LINE density in mammals (b) and LTR density in Plantae (c) showing generally good agreement between the two estimates. (d)-(f) Examples of comparisons of repeat density estimates between species. In each case we show box and whisker plots of the Pearson correlation of that density estimate with the Hi-C PC1 across chromosomes as well as the Pearson correlation between estimates.

